# Computationally engineered cyclic peptides reduce prion levels *in vitro*

**DOI:** 10.64898/2026.07.03.736251

**Authors:** Elpiniki Paspali, Carlos Oueslati Morales, Daria De Raffele, Adriano Aguzzi, Amedeo Caflisch, Simone Hornemann, Ioana M. Ilie

**Affiliations:** van ’t Hoff Institute for Molecular Sciences, University of Amsterdam, Amsterdam, The Netherlands; Swammerdam Institute for Life Sciences, University of Amsterdam, Amsterdam, The Netherlands; University of Zurich, Zurich, Switzerland; Institute for the Science of the Aging Brain (ISAB), St. Gallen Switzerland; Department of Biochemistry, University of Zurich, Zurich, Switzerland; Amsterdam Center for Multiscale Modeling (ACMM), University of Amsterdam, The Netherlands; Computational Soft Matter (CSM), University of Amsterdam, The Netherlands

## Abstract

Prion diseases are neurodegenerative disorders associated with the structural conversion of the cellular prion protein (PrP^c^) into its misfolded infectious isoform (PrP^Sc^). Despite substantial efforts, no disease-modifying therapy or cure is currently available. Here, we present an integrated computational-experimental pipeline for the rational design of cyclic peptides targeting PrP^c^ to inhibit its pathogenic conversion. Starting from crystal structures of antibody-bound mouse PrP^c^, we develop a rational design strategy combined with iterative molecular dynamics simulations and sequence optimization to generate peptides with enhanced binding and structural impact. Three candidates were selected for experimental validation. Our results show that 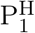 (^49^YGPDPSDSYT^58^, antibody numbering) that binds stably to the *α*_2_–*α*_3_ interface most effectively reduced PrP^Sc^ levels in GT1-7 cells, essentially by inducing allosteric re-arrangements that reinforce the intramolecular helical bundle. 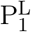 (^89^GQSNTKPYT^97^) and 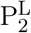 (^89^RQSNTWPYT^97^) binding the *β*_1_-*α*_1_/*α*_3_ junction exerted more modest effects due to the potential competition of the flexible tail to bind at this site. These results establish a mechanistic link between peptide-induced stabilization of PrP^c^ and inhibition of prion propagation and provide a generalizable framework for designing conformational stabilizers of aggregation-prone proteins.

## 1 Introduction

Prion diseases are fatal transmissible neurodegenerative disorders characterized by neuronal loss, vacuolation and glial activation and associated with the structural conversion of the cellular prion protein (PrP^c^) into its pathogenic scrapie isoform (PrP^Sc^).^1–3^ Unlike conventional proteinopathies, prion diseases are defined by a self-propagating mechanism in which the mis-folded conformer superimposes its aberrant structure onto native PrP^c^, thereby amplifying pathogenic assemblies and driving progressive neurodegeneration. ^4,5^ This mechanism drives disease progression by allowing PrP^Sc^ to multiply and disseminate from cell to cell, between brain regions and across individuals.

Structurally, native PrP^c^ consists of an intrinsically disordered N-terminal segment (residues 23–120) and a C-terminal globular domain (residues 121–231).^6–8^ The latter folds into three *α*-helices, *α*_1_, *α*_2_ and *α*_3_, and a short antiparallel *β*-sheet formed by *β*_1_ (128–131) and *β*_2_ (161–164) (numbering according to human PrP^c^).^9,10^ In contrast, PrP^Sc^ is enriched in *β*-sheet structure, is highly insoluble, resistant to proteolytic degradation ^11^ and potential therapeutic interventions.^12,13^ Given the different structural features associated with distinct biological responses, stabilizing the native conformation of PrP^c^, implicitly preventing its conversion to PrP^Sc^, represents an attractive therapeutic strategy.^12,14^

To date, no effective therapy has been developed to prevent the onset or progression of prion diseases. Despite extensive preclinical and clinical efforts, including approaches based on small molecules, *β*-sheet breaker compounds, and antibodies, no disease-modifying therapy has yet demonstrated efficacy in clinical trials. ^15,16^ For instance, Fe(III)-TMPyP, a small molecule binding a pocket formed by *α*_3_ and *β*_1_, has been shown to stabilize the native PrP^c^ conformation and reduce its susceptibility to pathogenic conversion as demonstrated in a protein misfolding cyclic amplification assay, prion-infected cells and prion-infected organotypic brain slices.^12,17,18^ Furthermore, structure-based virtual screening combined with biophysical validation has identified small molecules that bind native PrP^c^ with micromolar affinity and inhibit prion fibrillation.^19^ Beyond small molecules, antibodies have also emerged as promising therapeutic agents, with the anti-prion monoclonal antibody PRN100 providing early clinical evidence that direct target engagement in human prion disease is feasible. ^20^ Additionally, the monoclonal antibodies SAF34 and SAF61 have been shown to suppress PrP^Sc^ accumulation in prion-infected cells by promoting the accelerated degradation of PrP^c^.^21^ The POM antibody family further highlights the strong epitope dependence of PrP^c^ targeting.^22^ While POM1 binds to the globular domain and can induce neurotoxicity, POM2 recognizes the flexible N-terminal tail and induces lysosomal degradation of PrP^c^, thereby exerting neuroprotective effects, including suppression of POM1-induced neurotoxicity.^23–25^

In its monomeric state, PrP^c^ adopts a well-defined secondary structure in its C-terminal domain, yet, its surface lacks an accessible druggable site, a limitation for both small-molecule inhibitors and antibody-based strategies.^26,27^ Peptides, particularly cyclic peptides, overcome this challenge as they can be engineered to combine conformational rigidity and solubility, enabling high affinity binding to otherwise undruggable interfaces.^28–30^ Furthermore, cyclic peptides exhibit long in vivo stability, while maintaining robust antibody-like binding affinity and specificity.^28,29,31,32^ Over the past 25 years, more than 60 cyclic peptide-based therapeutics have received FDA and EMA approval, primarily as antibiotics, anticancer therapeutics, antifungals, antibacterials and immunomodulators. ^28,33–35^ These clinical advances position cyclic peptides as an attractive strategy for challenging targets such as PrP^c^, where their conformational rigidity and enhanced binding may enable stabilization of the native protein conformation and thereby prevent conversion into the pathogenic isoform.

Here, we pioneer the use of cyclic peptides as modulators of prion protein misfolding. We introduce an integrated framework for developing conformation-stabilizing cyclic peptide binders against the PrP^c^ by combining rational design, molecular dynamics simulations, iterative sequence engineering and experimental validation. More broadly, this work introduces a generic computational pipeline for engineering cyclic peptides against difficult-to-drug proteins, with applications beyond prions. Additionally, it provides a structural basis for exploring novel peptide design approaches.

## 2 Results

Cyclic peptides were designed and optimized through an iterative computational screening process prior to experimental testing (Fig. 1a). First, peptide sequences were rationally designed starting from the crystal structure of mouse PrP^c^ in complex with the POM1 antibody.^23^ Residues contributing most strongly to target binding were identified and grafted into the peptide sequence, which was subsequently cyclized in a head-to-tail manner, amino-to-carboxyl (Fig. S1). Second, extended molecular dynamics simulations of the resulting PrP^c^-peptide complexes were carried out to evaluate the stability of the interface. Third, the peptides were optimized through targeted single-point mutations to enhance solubility and interface properties (Fig. S1, Table S1). Fourth, the peptides were ranked according to their binding free energies calculated from umbrella sampling simulations. Finally, the top three ranked peptides were selected for experimental validation and in depth structural analysis.

**Figure 1.**
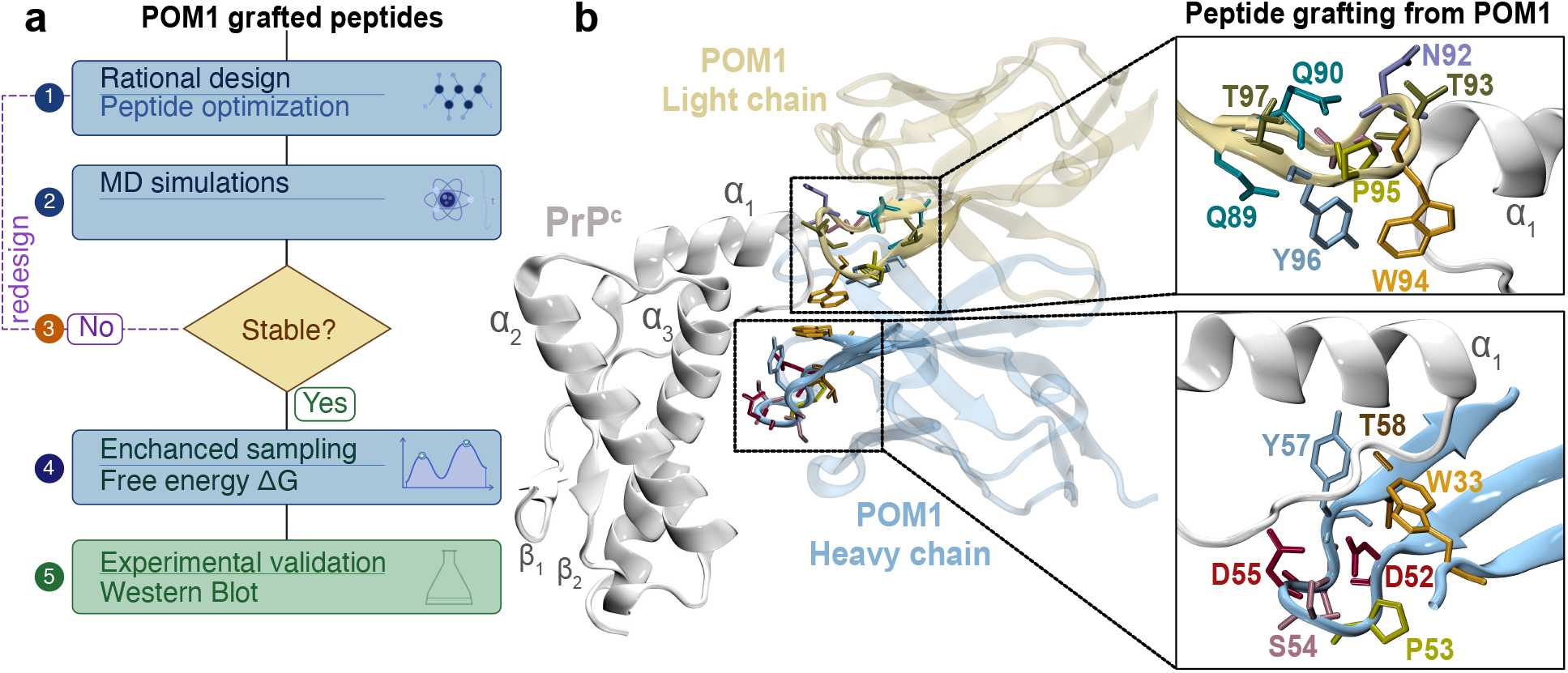
Peptide engineering pipeline. (a) Pipeline overview, (1) rational design and sequence optimization via single-point mutations, (2) molecular dynamics simulations to (3) assess complex stability via structural analysis (unstable designs were iteratively refined through the redesign cycle), (4) enhanced sampling to estimate the binding free energies (Δ*G*), (5) experimental validation of top candidates by Western blot. (b) Crystal structure of mouse PrP^c^ in complex with the POM1 monoclonal antibody (PDB: 4H88^23^). Key contact residues from the light and heavy chains were grafted to generate the linear peptides P^L^ and P^H^ (Fig. S1), which were subsequently head-to-tail cyclized and optimized.

### 2.1 Peptide design

The crystal structure of mouse PrP^c^ in complex with the monoclonal antibody POM1 was used as starting structure for this study (PDB ID: 4H88^23^) (Fig. 1b). To rationally design the peptides, we zoomed in on the PrP^c^-POM1 interface, focusing on the residues experimentally identified as critical for complex stability,^24^ *i*.*e*., W33, D52 and Y57, and Y50, S91, W94 and Y96 in the antibody heavy and light chains, respectively (Fig. 1b).

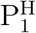 was derived from the heavy chain of the POM1 antibody, which binds the *α*_1_-*α*_2_ region of PrP^c^ (Fig. 1b). Specifically, residues ^52^DPSDSYT^58^ were first grafted from the POM1 heavy chain. Additionally, the experimentally identified critical residue W33 ^24^ was incorporated together with the common loop-forming residues glycine and proline, yielding the head-to-tail cyclized, renumbered sequence ^49^WGPDPSDSYT^58^ (Fig. S1a). The PrP^c^-peptide complex was subjected to 500-ns long all-atom molecular dynamics simulations to probe its stability (see Materials and Methods). During the first few nanoseconds (ns), the peptide rapidly detached from the PrP^c^ surface and reattached at the *α*_2_-*α*_3_ loop, where it remained stably bound for the remaining simulation (Fig. S1a). Upon binding, the peptide rearranged and the initially considered critical residues for binding, *i*.*e*., W33 became solvent exposed. Hence, to improve peptide solubility at the new binding site, the tryptophan residue was mutated to a tyrosine, generating a more hydrophilic variant while preserving the aromatic character of the peptide, resulting in the ^49^YGPDPSDSYT^58^ sequence Fig. S1a.

Peptides 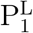 and 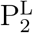 were derived from the POM1 light chain (Fig. S1b) starting from the isolated ^89^QQSNTWPYT^97^ sequence. To enable head-to-tail cyclization, the N-terminal glutamine was substituted with glycine, followed by ring closure to generate a conformationally constrained cyclic peptide. During the subsequent molecular dynamics trajectory, the peptide detached from its initial contact region on the PrP^c^ surface within the first 5 ns. After approximately 40 ns, the peptide reattached at *β*_1_-*α*_1_ loop, with occasional contacts formed with Q211 of the PrP^c^ *α*_3_ (Fig. S1b) and remained stably associated for the remaining of the simulation. Interestingly, the new binding site overlaps with the experimentally^36^ and computationally characterized POM1 epitope on PrP^c^,^26,27^ suggesting a potential competitive interference with the antibody binding. To improve solubility and hydrophilicity while maintaining the new binding site, the tryptophan residue was mutated to a lysine, yielding peptide 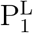, ^89^GQSNTKPYT^97^ (Table S1). Following the same protocol, 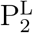 was generated by substituting the first glutamine with arginine to increase the local positive charge and enhance electrostatic complementarity at the binding interface, *i*.*e*., ^89^RQSNTWPYT^97^. Additionally, we designed the negative control peptide P^N^ (SNTWPYG) from the 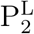 sequence through truncation and targeted mutations designed to disrupt hydrogen bonding networks and/or introduce steric clashes. This peptide detached from the PrP^c^ surface in all simulations and did not reattach throughout the simulations.

### 2.2 Binding stability and lead selection

Starting from the reattached PrP^c^-peptide configurations, five independent simulations were carried out for each protein-peptide complex for a cumulated sampling of 1.5 *µ*s to further assess the stability of the complexes. Analysis of the peptide displacements from the stable binding site showed that the peptides remained attached to PrP^c^ at this location in four of five runs for 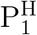 (deviations of 3.2 ± 1.8 Å (Fig. S2 a-c)), in all runs for 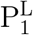 (deviations of 0.6 ± 0.03 Å (Fig. S2 d-f)) and three of five runs for 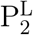 (deviations of 2.3 ± 1.1 Å (Fig. S2 g-i)). Note that in the 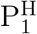 and 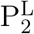 systems, the peptides only rearranged on the surface of PrP^c^ without detachment.

Next, we extracted representative structures of the PrP^c^-peptide complexes (via principal component analysis on the stable molecular dynamics trajectories (Fig. S3)) to be used as initial configurations for the enhanced sampling simulations (snapshots in Fig. 2a,b,c). We then performed umbrella sampling (US) simulations in triplicate for each PrP^c^-peptide complex, using the center-of-mass distance between the protein and peptide as the reaction coordinate. From the resulting free energy profiles (Fig. 2d), the binding free energies were calculated using 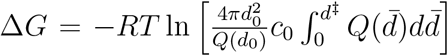.^37–39^ Here *d*_0_ is a distance beyond the interaction range, *d*^*‡*^ the location of the transition state of the binding reaction set to 2.5 nm, *R* is the gas constant and *c*_0_ is a reference concentration (typically 1 M). The distance-dependent partition function *Q*(*d*) was determined from the free energy profile, A(d) via *Q*(*d*) = exp[−*A*(*d*)*/*(*RT*)] at short distances and extrapolated beyond the interaction range using the entropy-dominated expression 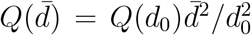. This calculation yielded −31 ± 1.1 kJ/mol, −24.5 ± 2.2 kJ/mol and −31 ± 3.4 kJ/mol for 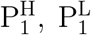 and 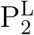, respectively (Fig. 2), which were selected as the top three candidates for experimental testing.

**Figure 2.**
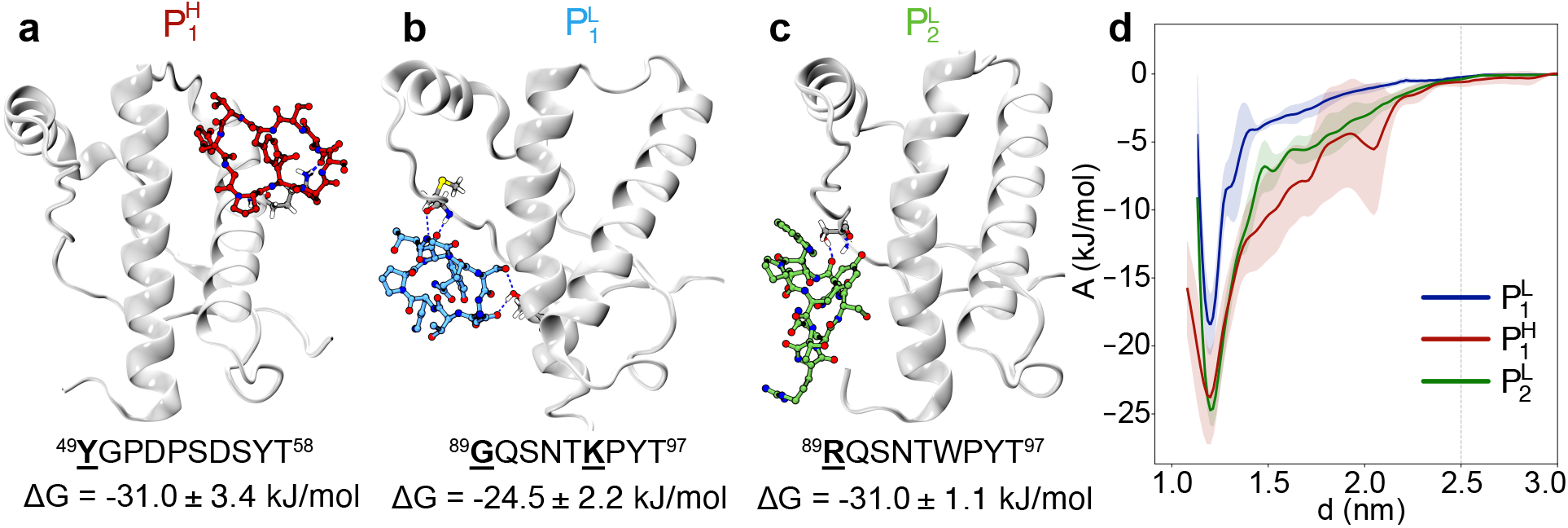
Top three ranked peptides. (a-c) Shown are representative structures of the stable PrP^c^-peptide complexes, the corresponding sequences and the calculated binding free energies. The underlined residues represent the modifications introduced to optimize binding. (d) Free energy profiles as a function of the center of mass distances between PrP^c^ and the peptides. The red, blue and green lines correspond to the profiles of 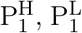 and 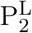, respectively. The shaded areas represent the error around the mean, calculated as the standard error of the mean from three independent replicates for each system

### 2.3 Peptides reduced PrP^Sc^ levels *in vitro*

The effect of the peptides on PrP^Sc^ levels were then investigated *in vitro*. We used the GT1-7 RML6 cell line, a persistently infected subline of immortalized mouse hypothalamic GT1-7 neuronal cells that stably propagates mouse-adapted RML6 (Rocky Mountain Laboratories, passage 6) prions. ^40^ Cells were treated for 90 h with each of the three peptides individually at a final concentration of 100 *µ*M. To maintain a constant peptide concentration throughout the experiment, the peptide-containing medium was replaced every 24h, resulting in three medium exchanges over the course of the 90 h treatment. The cells treated with the negative control peptide P^N^ (only performed in two of the experimental replicates due to limited peptide availability) or vehicle-only (PBS - VH1, VH2) were used as controls. Following treatment, the cells were lysed and an aliquot of each cell lysate was digested with Proteinase K (PK). Because PK selectively degrades the cellular prion protein PrP^c^ while leaving a protease-resistant core for PrP^Sc^, PK resistance is widely used as a surrogate marker of prion disease and enables the selective detection of PrP^Sc^.^41^ The total PrP levels (PrP^c^ and PrP^Sc^) and PK-resistant PrP^Sc^ levels were subsequently analyzed by Western Blotting (WB) and densitometry in non-digested and PK-digested samples, respectively, for each of the independent experimental replicates (Fig. 3 and Fig. S4).

**Figure 3.**
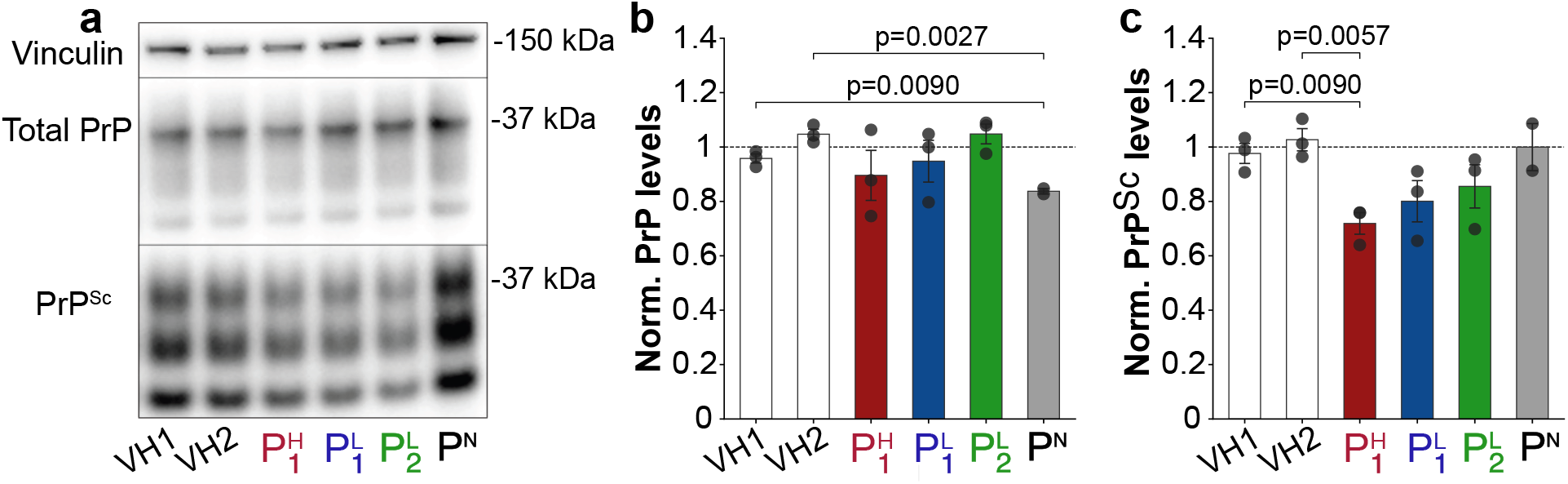
Effects of the peptides on total PrP and PrP^Sc^ levels. (a) WB images displaying total and PK-resistant PrP levels in GT1-7 cells following treatment with the peptides or vehicle (VH1, VH2). Vinculin served as a loading control. POM1 was used as detection antibody. kDa: kilo Dalton. (b, c) Quantification of the data shown in (a) and Fig. S4. using densitometry analysis. Data were normalized to the average of untreated controls and are presented as a scatter plot with bar. Shown are the mean and the error bars of the standard deviation. Each data point represents an independent biological replicate (n=3 cultures for the vehicle controls and peptides; n=2 for P^N^). The statistical significance was assessed using Brown–Forsythe and Welch ANOVA test, P < 0.05.

The WB analysis of non-PK-digested samples revealed marginal differences after peptide treatment (middle panel in Fig. 3a). This observation is supported by the quantitative densitometry analysis, which showed that cells treated with the peptides (including the negative control P^N^) and untreated cells exhibited PrP levels comparable to those of the vehicle control (VH1 baseline) (Fig. 3b). These findings indicate that the peptide treatment does not alter overall PrP^c^ expression. In contrast, the analysis of PK-digested samples showed a clear reduction in signal intensity in peptide-treated cells, indicating lower levels of PrP^Sc^ (bottom panel in Fig. 3a). Importantly, this reduction cannot be attributed to altered expression levels of PrP^c^, as treatment of GT1-7 RML6 cells with 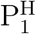 did not significantly affect total PrP expression levels compared to vehicle-treated cells (Fig. 3b). Among the tested peptides, 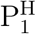 had the strongest effect, reducing PrP^Sc^ levels by approximately 30% compared with the vehicle control. 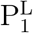 and 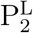 induced more moderate effects (approximately 0.8-fold), but these effects did not reached statistically significance. In contrast, the negative control peptide and untreated cells showed PrP^Sc^ levels comparable to those of vehicle-treated cells. These results demonstrate that 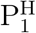 selectively and significantly reduces PrP^Sc^ levels without affecting total PrP levels.

### 2.4 Multivalent interaction networks stabilize the complex interface

The analysis focused on the interfacial interactions revealing that 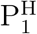 (^49^YGPDPSDSYT^58^) binds primarily the *α*_2_-*α*_3_ interface of PrP^c^ (Fig. 2a), with the main stabilizing interaction being a salt bridge between 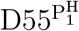 and 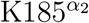 Fig. 4a, 67% occupancy). Additionally, transient hydrophobic contacts were formed by 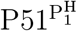 and 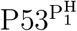 with 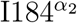 (32% and 34%), 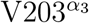 (34%), 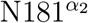 (28%) forming a hydrophobic cluster encompassing *α*_2_ and *α*_3_ on the PrP^c^ surface (Fig. 4a right snapshot). Thus, the favorable binding free energy of Δ*G* = −31.0 ±3.4 kJ/mol (Fig. 2d, Fig. 4a) arises mainly from the electrostatic contribution of the 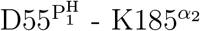 salt bridge.

**Figure 4.**
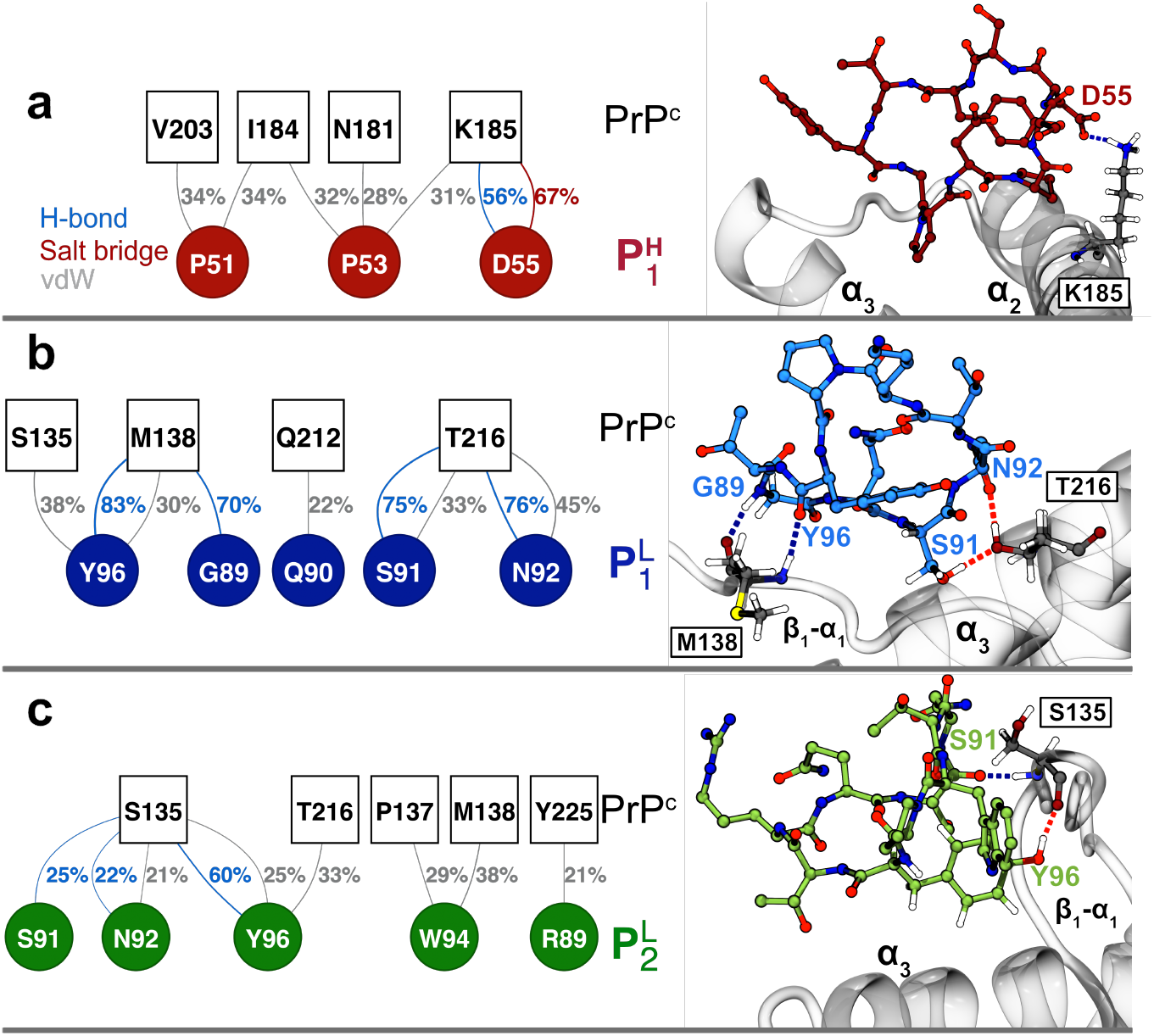
Intermolecular contacts of the PrP^c^-peptide complexes. Shown are (left) contact maps of the intermolecular interactions between 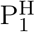 (a, red), 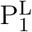 (b, blue) and 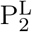 (c, green) with PrP^c^, and (right) representative peptide binding poses. PrP^c^ is shown in cartoon representation and the peptides as sticks.

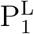 (^89^GQSNTKPYT^97^) binds the *β*_1_-*α*_1_/*α*_3_ junction through hydrogen bonds. Specifically, 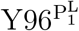 forms a persistent backbone hydrogen bond with 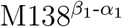 (83% occupancy), anchoring the peptide to the *β*_1_-*α*_1_ loop (Fig. 4b). Additionally, 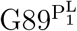 engages 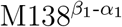 from the opposite direction through a backbone hydrogen bond (70% occupancy). On the *α*_3_ helix, 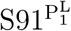 and 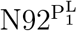 both contact 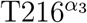 through sidechain hydrogen bonds (75% occupancy), establishing a second anchoring point that bridges the peptide to the C-terminal helix. The W94K substitution introduced in the optimization cycle eliminates the bulky indole side chain present in the parent sequence (P^L^), however, the introduced lysine at position 94 did not form long-lived intermolecular contacts, indicating that this mutation reduces steric constraints rather than contributing new favorable interactions. These interactions reflect in the resulting potential mean forces extracted from the umbrella sampling simulations, which yielded a binding free energy of Δ*G* = −24.5 ± 2.2 kJ/mol (Fig. 2b).

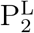 (^89^RQSNTWPYT^97^) retains the native tryptophan at position 94 and carries the Q89R substitution, introducing a positively charged arginine at the N-terminus (Fig. 4c). Upon interaction with the protein, Y96 serves as the primary anchor, forming a stable hydrogen bond with 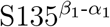 (60% occupancy) and transient contacts with 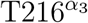, 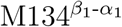, 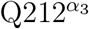, and 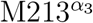 and a *π*-*π* stacking interaction with Y226^C-term^ positioning it at the *β*_1_-*α*_1_ loop and *α*_3_ helix. Essentially, 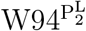 forms a network rich in aromatic and polar interactions with 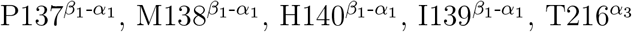, and 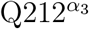 (unlike 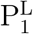, where the W94K substitution removed the indole ring). The introduced 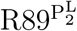 forms transient cation-*π* interactions with Y225^C-term^, extending the contact toward the C-terminus of *α*_3_. 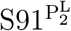 and 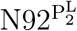 forms hydrogen bonds with 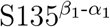 (25% and 22%, respectively).This network of contacts yielded Δ*G* = −31.0 ± 1.1 kJ/mol (Fig. 2c), comparable to 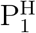.

### 2.5 Peptide binding remodels the intramolecular contact network

The analysis focused on the protein plasticity revealed that all three peptides consistently reduced PrP^c^ flexibility at the *β*_2_-*α*_2_ loop (residues 166-172, Fig. 5a) independently of their binding site. This shared pattern independently of the peptide binding site suggests that the observed effect is not driven by direct peptide-loop interactions, but rather by indirect stabilization propagated through the helical bundle. Notably, the *β*_2_-*α*_2_ loop is dynamic^26,42,43^ and amino acid substitutions within this region were shown to modify local structural rigidity^42,43^ and thereby impact the transmission efficiency of prion diseases. ^44,45^ In contrast, the effect on the *β*_1_-*α*_1_ loop is system-dependent with 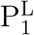 reducing the flexibility of this loop relative to free PrP^c^, whereas 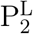 has marginal impact despite binding at the same *β*_1_-*α*_1_/*α*_3_ junction. Interestingly, 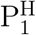, which contacts the *α*_2_-*α*_3_ interface, increases *β*_1_-*α*_1_ loop flexibility in an allosteric manner.

**Figure 5.**
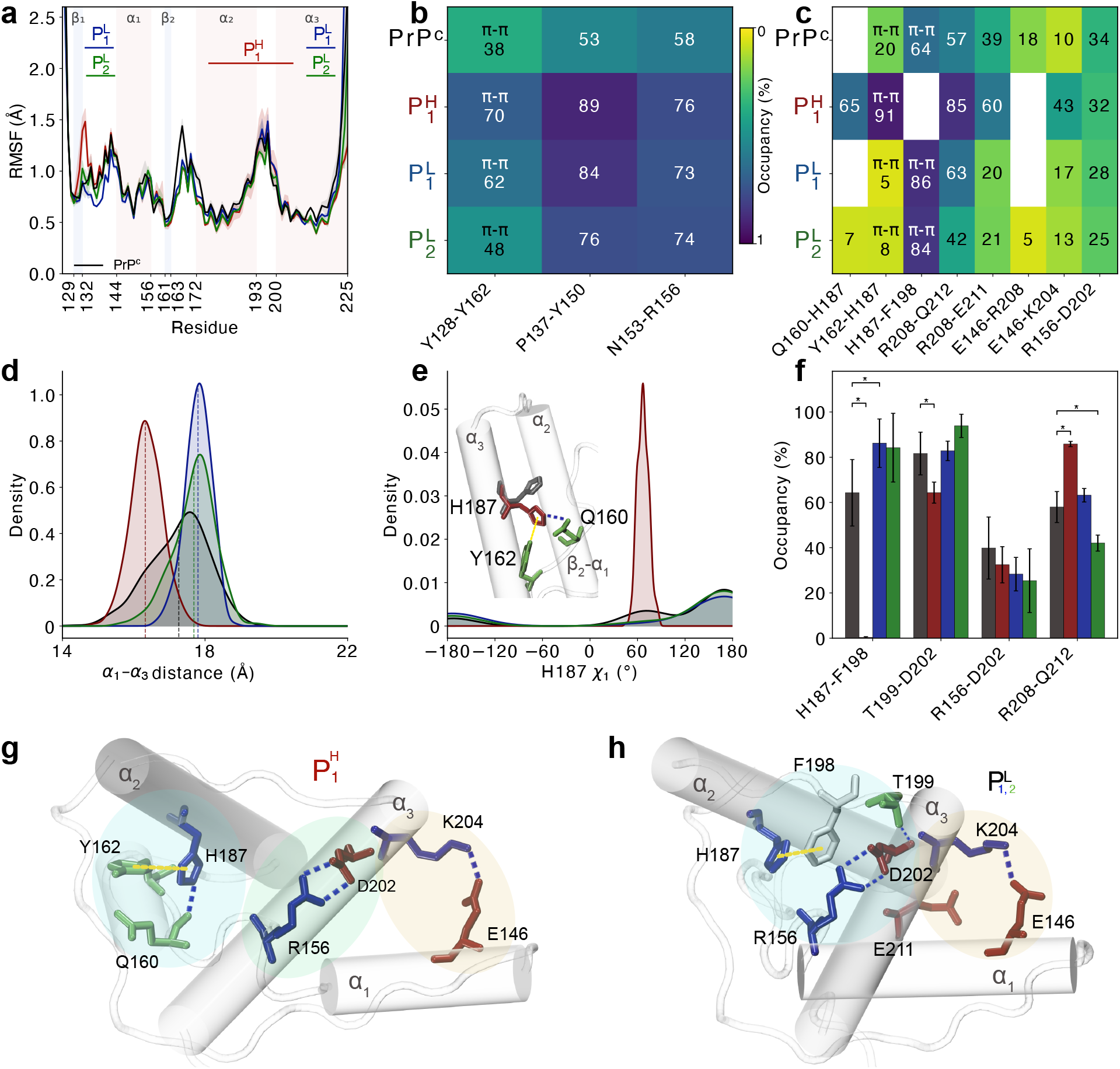
Peptide binding modulates PrP^c^ backbone dynamics and intramolecular contact network. **a)** Root mean square fluctuations (RMSF) profiles of PrP^c^ (black), 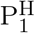-bound (red), 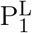-bound (blue), and 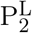-bound (green) states averaged over independent 100-ns windows from trajectories with a stable interface Fig. S2). Shaded areas: standard error. Secondary structure elements and peptide binding sites are indicated. **b–c)** Intramolecular contact heatmaps of PrP^c^ the free state and peptide–binding states. Each cell shows the mean occupancy (%) and the dominant interaction type (*π*–*π* stacking; unlabeled cells indicate hydrogen bonds). **d)** Kernel density estimate (KDE, bandwidth = 0.35) of the *α*_1_–*α*_3_ inter-helix distance (residues 144–156 and 200–225, C*α* centres of geometry. Dashed lines: system means. **e)** KDE (bandwidth = 0.25) of the H187 *χ*_1_ dihedral (N–C*α*–C*β*–C*γ*) in apo PrP^c^ (gray) and peptide-bound states (red:P^H^). Inset: representative structures of the dominant H187 rotamers. **f)** Intermolecular contact occupancies (%). Error bars: SEM across independent runs. Brackets denote significant differences from the free state (Mann–Whitney *U* test; * *p <* 0.05, ** *p <* 0.01, *** *p <* 0.001). **g–h)** Representative anchoring snapshots. 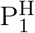 (g) contacts *β*_2_–*α*_3_ via hydrogen bonds and *π*–*π* interactions, and bridges *α*_1_–*α*_3_ through R156–D202 and E146–K204 salt bridges. 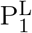 and 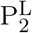 (h) share the R156–D202 and E146–K204 salt bridges, with H187, F198, and T199 forming additional hydrogen bonds with D202.

To further understand the driving interactions that modulate the plasticity of the loops, we analyzed the intramolecular contacts within PrP^c^ that were altered upon peptide binding relative to the free protein (Fig. 5b,c). Three contacts were consistently strengthened across all three peptide-bound systems, albeit with a less significant effect in 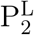 (Fig. 5b). First, the *π*-*π* stacking interaction between 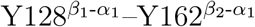 bridging the two *β*-strands, increased from 38% to 70, 62 and 48% for 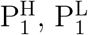 and 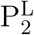, respectively. Second, the 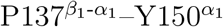 hydrogen bond connecting the *β*_1_-*α*_1_ loop and *α*_1_ was strengthened from 53% in the free state to 76-89% across all peptide-bound systems. Third, the 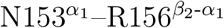 hydrogen bond between *α*_1_ and *β*_2_-*α*_1_ was reinforced from 58% to 73-76%. Together, these shared rearrangements indicate that peptide binding consistently stabilizes the *β*-sheet scaffold and its contacts with *α*_1_ as indicated by the stable inter-*β*-strand distances in all systems ((Fig. 5d)). Beyond these shared changes, peptide binding induced a distinct pattern of contact re-organization. Upon 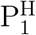 binding, two new contacts bridging the *β*_2_-*α*_2_ junction were formed, 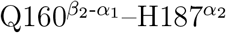 (65%) and 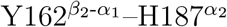 (91%), which contribute to the reduced flexibility of the *β*_2_-*α*_2_ loop (Fig. 5c). These new interactions redirect the interhelical contacts of *α*_3_ with the *α*_2_–*α*_3_ loop to contacts with the *α*_1_–*β*_2_ loop, reflected in the reduced distances between the *β*_1_-*α*_1_ and *β*_2_-*α*_2_-loops and between *α*_1_ and *α*_3_ (Fig. 5d) relative to the free state. Additionally, the new contact formation is accompanied by the abolition of the H187– F198 contact (64% to 0%, Fig. 5c), which requires a different 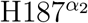 rotamer (Fig. 5d, (Fig. 5e). Specifically, the *χ*1 dihedral analysis shows that 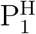 locks H187 in the trans rotamer (*χ*1 ≈ −67° ± 7°, Fig. 5e, redirecting the imidazole ring toward 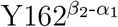 (92% occupancy) and 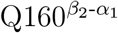 (66% occupancy) to the *β*_2_-*α*_2_ loop (Fig. 5c, d) This rotamer switch is structurally analogous to the effect of H187R Gerstmann–Sträussler–Scheinker syndrome mutation, which replaces 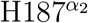 with a permanently charged residue and disrupts the same electrostatic network.^46,47^ However, the 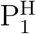-induced switch is accompanied by compensatory contacts (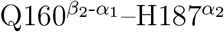 (65%) and 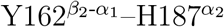 (91%)) that prevent the hydrophobic core exposure characteristic of the pathogenic mutation^48^ (Fig. 5c). P^H^ also stabilizes the 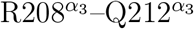 bond (from 57% to 85%), the 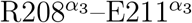 intrahelical bond (60%) and the 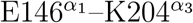 (43%) interhelical salt bridge while preserving the 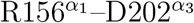 interaction (Fig. 5f, g). Together, this interaction triad bridges *α*_1_ and *α*_3_ at two distinct sites instead of one in the free state (Fig. 5g), resulting in a tighter bundle of the two helices (Fig. 5d).

Upon 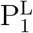 and 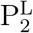 binding, the 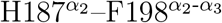 *π*-*π* interaction, which connects the *α*_2_ helix with the *α*_2_-*α*_3_ loop is strengthened from 64% to 86% and 84%, respectively, relative to free PrP^c^ (*χ*_1_ ≈ 180)(Fig. 5c), leading to H187 retaining the PrP^c^ rotamer state (Fig. 5e) and forming contacts with the *α*_2_–*α*_3_ loop though F198 (Fig. 5c, h), in contrast to P^H^ which locks H187 into an alternative rotamer (*χ*_1_ ≈ 60) engaging the *β*_2_–*α*_1_ loop, resulting in a more compact helical bundle as reflected in the reduced *α*_1_–*α*_3_ distance (Fig. 5d). Furthermore, the *α*_2_-*α*_3_ and *α*_1_ connection is maintained through the 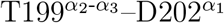 hydrogen bond (Fig. 5c). The latter also retains the native salt bridge with R156 and together with the reinforcement of the 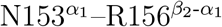 bond (Fig. 5f) has a comparable effect as the attachment of the POM1 and POM6 monoclonal antibodies.^26^

### 2.6 Relationship to the pathologic R208 mutation

The R208H mutation is linked to familial Creutzfeldt-Jakob disease and was proposed to destabilize PrP^c^ by perturbing electrostatic interactions and backbone dynamics within *α*_3_.^49^ Our simulations show that in free PrP^c^, 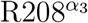 anchors the *α*_1_-*α*_3_ interface through an alternating salt bridge between 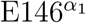 and participates in an intra-*α*_3_ hydrogen bond with 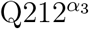 (57%) (Fig. 5c, f). Upon peptide binding, the 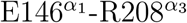 salt bridge was reduced for all three peptides. Specifically, 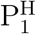 binding enhanced the 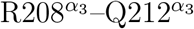 hydrogen bond (Fig. 5(f)) and the 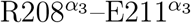 salt bridge (Fig. 5(c)) (from 57 to 85% and from 26 to 40%, respectively). The 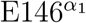 switched partner to 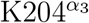, forming a new *α*_1_-*α*_3_ salt bridge (10 to 43%) that functionally replaced the lost 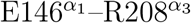 interaction (Fig. 5(c, f). Additionally, the 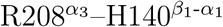 hydrogen bond previously identified as a key conformational switch induced by the neurotoxic POM1 antibody^24^ did not form in any of our peptide-bound systems, distinguishing the peptide-induced perturbations from the neurotoxic POM1-induced rearrangement.

## 3 Discussion

Monoclonal antibodies have been developed to recognize epitopes along the PrP^c^ sequence^21,22^ and to act on PrP^c^ to prevent its transition into the *β*-sheet–rich PrP^Sc^ isoform and the downstream cascade leading to prion diseases. ^50–52^ However, their therapeutic application remains limited by poor blood–brain barrier penetration, heterogeneous distribution, and safety concerns,^53^ as certain anti-PrP antibodies can induce neurotoxic effects.^23,54^ In contrast, cyclic peptides represent a promising alternative, due to their enhanced structural stability, resistance to proteolytic degradation and synthetic accessibility, while enabling robust antibody-like target binding. ^28,29,37^ Here, we present a peptide design pipeline that integrates rational design, iterative molecular engineering, *in silico* validation and *in vitro* testing to generate cyclic peptides that bind to PrP^c^ and reduce PrP^Sc^ levels. These peptides were designed to stabilize the native PrP^c^ conformation, thereby inhibiting its conversion into the pathogenic PrP^Sc^ isoform and blocking its subsequent misfolding and aggregation. Their cyclic architecture may also offer favorable properties for further optimization, including improved stability and delivery.^29^

The three cyclic peptides were designed based on the heavy and light chain of the POM1 monoclonal antibody, 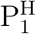 (^49^YGPDPSDSYT^58^), 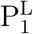 (^89^GQSNTKPYT^97^) and 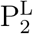 (^89^RQSNTWPYT^97^) that bind PrP^c^ at two distinct surface regions of PrP^c^, both of which are functionally linked to prion conversion and thus represent attractive sites for therapeutic intervention. 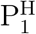 binds the *α*_2_-*α*_3_ helical interface, partially overlapping with the epitope recognized by POM1 (*α*_1_-*α*_3_) and shared with the POM6 and POM7 (*α*_1_-*α*_2_) monoclonal antibodies.^22^ These monoclonal antibodies elicit distinct biological responses. POM1 binding is associated with neurotoxicity that is independent from prion infectivity and replication, whereas POM6 binding promotes a non-toxic conformational ensemble. ^22–24^

Our data show that 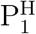 significantly reduced PrP^Sc^ levels in prion-infected GT1-7 cells, demonstrating anti-prion activity in a biologically relevant *in vitro* model and identifying the *α*_2_-*α*_3_ interface as a functionally important site for modulating prion conversion. Supporting this interpretation, molecular dynamics simulations revealed that 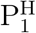 anchors at this interface through a persistent 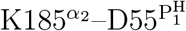 salt bridge, initiating an allosteric response that propagates throughout the globular domain and promotes conformational stabilization. This includes a 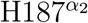 rotamer shift that enables new intradomain interactions within *β*_2_-*α*_1_ and promotes formation of an 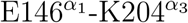 interhelical salt bridge, thereby tightening the *α*_1_–*α*_3_ bundle. These results suggest that 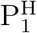 inhibits prion conversion by reinforcing native interdomain packing.

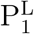 and 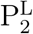 bind at the *β*_1_-*α*_1_ and *α*_3_ region, encompassing the segment of residues 106-147, which was identified as central to fibril formation.^55^ Specifically, small molecules that bind with micromolar affinity to the *β*_1_-*β*_2_/*α*_2_ region have been shown to inhibit prion formation by stabilizing the native conformation and blocking conformational conversion into PrP^Sc^.^19^ However, our cell-based prion-infectivity assay reveals that 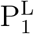 and 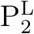, did not significantly reduce PrP^Sc^ levels in GT1-7 cells. Our molecular dynamics simulations offer a structural rationale for this outcome. Although both peptides bind the *β*_1_-*β*_2_/*α*_2_ region, they do not reinforce coupling between the *β*_1_–*α*_1_–*β*_2_ subdomain and the *α*_2_–*α*_3_ helical core. Specifically, 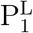 interacts with PrP^c^ primarily through backbone hydrogen bonds, while the introduced 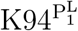 residue forms no persistent intermolecular contacts, resulting in relatively weak binding (Δ*G* = −24.5±2.2 kJ/mol). Moreover, unlike 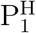, which stabilizes the 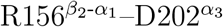 and 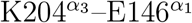 salt bridges, 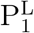 and 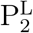 rely mainly on weaker hydrogen-bond-mediated anchoring at the *α*_2_-*α*_3_ interface. Consequently, they preserve interhelical distance distributions of the free PrP^c^, whereas 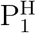 tightens the *α*_1_–*α*_3_ interface. Since detachment of the *β*_1_–*α*_1_–*β*_2_ subdomain from the helical core was shown to initiate PrP^c^ to PrP^Sc^ conversion,^56–58^ the stronger interdomain stabilization induced by 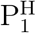 may explain its superior anti-prion activity. Additionally, our previous work showed that the flexible tail of PrP^c^ frequently interacts with the *β*_1_-*β*_2_/*α*_2_ region and that these contacts are altered by truncations or antibody binding.^27^ Since our experiments were performed using full-length prion protein, the overlapping binding sites of 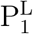 and 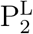 with this dynamically contacted region may further reduce their effect in the cellular environment.

## 4 Conclusion

We developed an iterative computational pipeline to rationally design cyclic peptides starting from the crystal structures of PrP^c^ in complex with the POM1 monoclonal antibody, followed by optimization through sequential rounds of molecular dynamics simulations, targeted single-point mutations and enhanced sampling. Using this technique, we selected three cyclic peptides, that showed the most favorable binding to PrP^c^, to advance into experimental validation. We identified 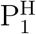 (^49^YGPDPSDSYT^58^) as the most potent PrP^c^ binder, which also achieved the strongest reduction of PrP^Sc^ levels in GT1-7 RML6 cells without affecting total PrP expression. These findings establish a mechanistic link between targeted PrP^c^ binding of peptides and impaired prion propagation, providing a peptide-based strategy to inhibit conversion into the scrapie isoform. More broadly, this engineering pipeline offers a platform for targeting aggregation-prone proteins beyond prion diseases.^28,37,59–61^

## 5 Materials and methods

### 5.1 System preparation

To develop cyclic peptides that compete against monoclonal antibodies against PrP^c^ binding, the 124-225 segment of the PrP^c^ was extracted from the crystal structure of mouse PrP^c^ in complex with the monoclonal antibody POM1^36^ (PDB ID: 4H88). The N- and C-termini of the protein were capped with acetyl and N-methylacetamide groups, respectively. All histidine side chains were modeled as neutral and protonated at N_*δ*_.

### 5.2 Simulation protocol

Three sets of simulations were carried out using the same simulation parameters. First, unrestrained 500 ns molecular dynamics simulations were carried out in the canonical ensemble to investigate the stability of the initially designed PrP^c^-peptide complexes. In all cases, the peptides detached from the surface of PrP^c^ and reattached at secondary locations. The new protein-peptide configurations were then advanced into further MD simulations. Mutations were introduced in the peptides to improve solubility or to enhance binding to the protein (see Section 2.1). Second, five 300-ns simulations of each selected complex were run to probe the stability of the complexes and characterize the PrP^c^-peptide interactions. All the independent simulations were initiated from identical starting conformations, with each replicate being assigned distinct initial velocities. Third, umbrella sampling simulations were performed to compute the binding free energies of the peptides to PrP^c^.

### 5.3 Molecular dynamics simulations

All simulations were carried out using the GROMACS 2020.4 simulation package. The simulations were performed using the all-atom CHARMM36m force field^62,63^ and the TIP3P water model.^64^ The systems were solvated in cubic boxes with edge lengths of 10 nm for the PrP^c^-peptide complexes. Each system was neutralized and a background concentration of 150 mM of NaCl was added. Steepest decent energy minimization was followed by a two-step equilibration, *i*.*e*., 5 ns NVT equilibration with position restraints of 1000 kJ·mol^−1^·nm^−2^ applied on the backbone atoms of the protein and peptide, followed by a 5 ns NPT equilibration in absence of restraints. The temperature and pressure were kept constant at of 300 K and 1 bar using the velocity rescaling^65^ (modified Berendsen) thermostat and Berendsen barostat,^66^ respectively. The temperature and pressure coupling times were fixed to 0.1 and 2 ps, respectively. The production simulations were performed in the NVT ensemble in absence of restraints for 300 ns per replica. The potential smoothly converges to zero at the cutoff using the Verlet cutoff scheme and the short-range interactions were cut off beyond 1.2 nm. The long-ranged electrostatic interactions were evaluated using the particle mesh Ewald (PME) with a cubic interpolation order, a real space cutoff of 1.2 nm and a grid spacing of 0.16 nm.^67,68^ The bond lengths were constrained by means of a fourth order LINCS algorithm with 2 iterations.^69^ In all simulations, the time step was fixed to 2 fs, and periodic boundary conditions were applied.

### 5.4 Umbrella sampling

To determine the binding free energies between the peptides and PrP^c^, umbrella sampling (US) simulations were performed.^70,71^ Briefly, a harmonic biasing potential was applied to restrict sampling to discrete windows along the center-of-mass distance, *d*, between the peptide and PrP^c^ (reaction coordinate). The windows were centered at values *d*_*i*_ where the harmonic bias potential *ω*_*i*_(*d*) only restricts the reaction coordinate in the *i* ^th^ window to fluctuate around *d*_*i*_ by:^72^ 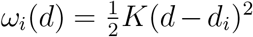, with *K* the force constant. The unbiased free energy for window *i, A*_*i*_(*d*), was extracted from the probability distribution 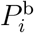 and the bias potential and was obtained by

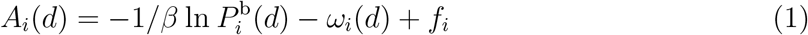

where *β* = 1*/k*_*B*_*T*, *k*_*B*_ being the Boltzmann constant, *T* the temperature and *f*_*i*_ a window-dependent offset. The windows were combined using the weighted histogram analysis method (WHAM)^73–75^ to determine *f*_*i*_.

The starting configurations of PrP^c^-peptide complexes were selected from stable MD trajectories. In preparation for the US simulations, the peptide was pulled away from the protein along the z-axis using a harmonic potential at a rate of 0.008 nm/ps for 400 ps (total COM displacement 3.15nm), with a force constant of *K* =1000 kJ/(mol nm^2^). Additionally, the backbone of the protein was restrained with a force constant of 1000 kJ/(mol nm^2^). Each window was first equilibrated in the NVT ensemble for 10 ns followed by 5 ns NPT equilibration, with position restraints applied on the backbone atoms of the protein and the peptide. For the umbrella sampling simulations, the simulation box was reshaped from the cubic geometry used in the unbiased MD runs (9.86 × 9.86 × 9.86 nm) to a rectangular box of 6.4 × 6.4 × 9.86 nm, with the longest edge aligned along the pulling direction (z). For the production three different replicated of each system were generated to probe convergence. In all cases, each umbrella window was simulated in the NPT ensemble for 200 ns.

### 5.5 Principal Component Analysis

To identify representative binding modes for each peptide–PrP^c^ complex, principal component analysis (PCA) was performed on the concatenated MD trajectories using the MD-Analysis PCA module.^76,77^ PCA was applied to the C*α* atoms of the peptide aligned on the PrP^c^ backbone, such that resulting components describe the repositioning of the peptide relative to the receptor. All replicas were concatenated prior to analysis to span a shared conformational space (Fig. S3). Conformational clusters were identified with HDBSCAN ^78^ applied to the first four principal components. Cluster centroids were defined as the trajectory frames closest to the center of mass of each cluster in PC space. The centroid structures were used as starting conformations for umbrella sampling simulations.

### 5.6 Residue interaction network analysis

Intermolecular contacts between each peptide and PrP^c^ were characterized using RING,^79^ which classifies residue-residue interactions (hydrogen bonds, salt bridges, hydrophobic contacts) based on geometric and physicochemical criteria under strict cuttofs (hydrogen bond distance: 3.5Å, angle: 150°). Frames were extracted per ns of the trajectories for each replica (300 frames per replica) and processed with RING. The interaction occupancy for each residue pair was computed as the fraction of frames in which the contact was observed, averaged across replicas. Per replica, the occupancies were then aggregated across the *n*_rep_ independent trajectories (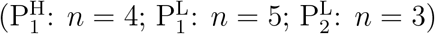 : *n* = 3), and the mean occupancy, standard deviation and standard error of the mean were reported. For the peptide binding analysis, only contacts with occupancy >20% were reported. To assess whether peptide binding perturbs the intramolecular contact network of PrP^c^, the same RING analysis was performed on apo PrP^c^ simulations. For each intramolecular residue pair, the per-replica occupancy difference Δ*θ* = *θ*_bound_ − *θ*_apo_ was computed across replicas. Statistical significance was assessed using the Mann–Whitney *U* test (two-sided),^80^ a non-parametric rank-sum test that makes no assumption of normality — appropriate given the small number of replicas. *p*-values were corrected for multiple comparisons using the Bonferroni procedure (*α*_corr_ = 0.05*/N*_pairs_), and only pairs with 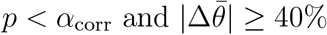 were considered significant.

### 5.7 Culture of GT1-7 and peptide treatment

GT1-7 cells^40^ were cultivated in sterile-filtered Opti-MEM medium (Gibco #31985070) supplemented with 10% FBS (Cytiva, SV30160.03), 1% Non-Essential Aminoacids (Gibco #11140050), 1% GlutaMAX (Gibco #35050061) and 1% Penicillin/Streptomycin (Gibco #15140122). The cells were kept under sterile conditions, in a humidified incubator at 37^°^C and 5% CO_2_. When the cells reached approximately 80% confluence, they were passaged 1:3-1:5 using Accutase (Gibco #A1110501).

Peptides P^N^, 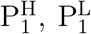 and 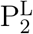 were purchased from Biosynth and received frozen in dry ice. The peptides were stored at −80°C and thawed slowly buried in wet ice, immediately before use. Afterwards, peptides were kept at 4 °C.

### 5.8 PK Western Blotting

Following the treatment with 100 *µ*M peptides, the cells were washed with sterile PBS (Gibco #10010023) before being lysed (Tris-HCl pH 8.0, 0.05 M, NaCl 0.15 M, Nadeoxycholate 0.5%, Triton X-100 0.5 % in ddH2O). Cell lysates were then centrifuged at 10,000 g for 10 min. A bicinchoninic acid (BCA) assay (Pierce, Thermo Fisher Scientific) was used to measure the total protein content of each sample. For immunoblotting, the samples were diluted to achieve the same total protein concentration, and, if required, digested using PK (final concentration 2.5 *µ*g/ml). Digestion was stopped with boiling the samples after addition of 1 mM final Dithiothreitol (DTT, Bio-Rad, Hercules, CA, USA) in NuPAGE 4× LDS loading buffer (Thermo Fisher Scientific). Subsequently, the samples were loaded onto 4-12% Bis-Tris NuPAGE gradient gels (Invitrogen, Thermo Fisher Scientific) and blotted onto a nitrocellulose membrane using the iBlot dry transfer system (Invitrogen, Thermo Fisher Scientific). The membranes were blocked using 5% SureBlock (LubioScience) diluted in 1×PBS containing 0.1% Tween-20 (PBST, Sigma Aldrich) for 30 min. The membranes were then incubated with the anti-PrP primary antibodies (POM1 (666 ng/ml), POM2 (1 *µ*g/ml)) diluted in CanGet solution 1 (Toyobo Research Reagents #NKB-101) or the anti-Vinculin primary antibody (#ab129002, 1:2000), diluted in 1% SureBlock-PBST, overnight at 4^°^C under shaking conditions. For detection, anti-mouse horseradish peroxidase (HRP) or anti-rabbit HRP (Bio-Rad) was diluted 1:5,000 in 1% SureBlock-PBST. The imaging was performed on LAS-3000 System (Fujifilm, Tokyo, Japan).

## Acknowledgements

I.M.I. acknowledges support from the Sectorplan Bèta & Techniek of the Dutch Government, the Dementia Research - Synapsis Foundation Switzerland and the Molecular Material Design Technology Impulse grant. S.H. acknowledges support from the Michael J. Fox Foundation (MJFF-020710, MJFF-021073), the Foundation for Research in Science and the Humanities at the University of Zurich (STWF-24-11) and the Innovation Fund of the University Hospital of Zurich (INOV00169). The computational resources were provided by the Dutch National Supercomputer Snellius. This work has been carried out within the framework of the COST Action CA2311-SNOOPY ‘Searching for Nanostructured or pOre fOrming Peptides for therapY’, supported by COST (European Cooperation in Science and Technology).

