## Supplementary information for "Computationally engineered cyclic peptides reduce prion levels *in vitro*"

### 1 Peptide design and properties

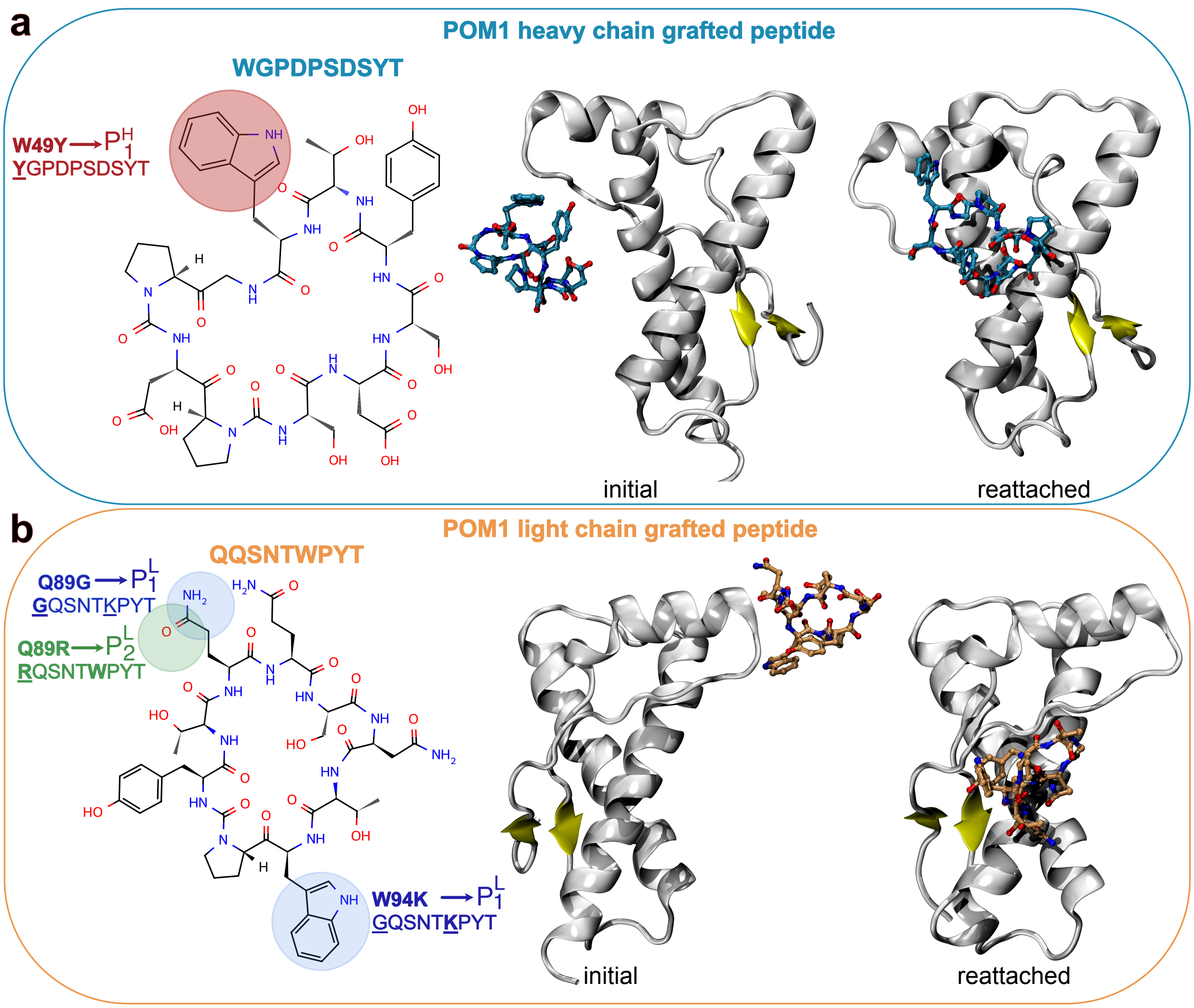


**Fig. S1 Grafted peptide systems derived from POM1 antibody. a)** POM1 heavy chain grafted peptide (WGPDPSDSYT) and its variant: $P^H_1$. **b)** POM1 light chain grafted peptide (QQSNTWPYT) and its two variants: $P^L_1$ and $P^L_2$. For each system, simulation snapshots at 1 ns and 500 ns are shown to highlight the initial (left snapshot) and the reattached site of the peptides relative to $PrP^c$ (white cartoon), with the peptide shown as sticks. Chemical structures of the parent peptides are shown on the left, with mutation sites circled.

**Table 1** Estimated water solubility of the cyclic peptides. Solubility was classified heuristically: peptides with >50% hydrophobic residues (V, I, L, F, W, M) were predicted as poorly soluble; those with ≥25% charged residues (D, E, K, R, H) as well soluble, alternative; the sign of the mean Hopp & Woods hydrophilicity score determined the classification.

| Peptide | Sequence | Residues | Charged (%) | Hydrophobic (%) | Avg hydrophilicity | Solubility |
|---|---|---|---|---|---|---|
| $P_1^L$ | GQSNTWKPYT | 9 | 11 | 0 | 0.07 | Good |
| $P_2^L$ | RQSNTWPYT | 9 | 11 | 22 | −0.31 | Poor |
| $P_1^H$ | YGPDPSDSYT | 10 | 20 | 0 | 0.16 | Good |
| $P^L$ | QQSNTWPYT | 9 | 0 | 22 | −0.62 | Poor |
| $P^H$ | WGPDPSDSYT | 10 | 20 | 10 | 0.05 | Good |

#### 2 Complex stability

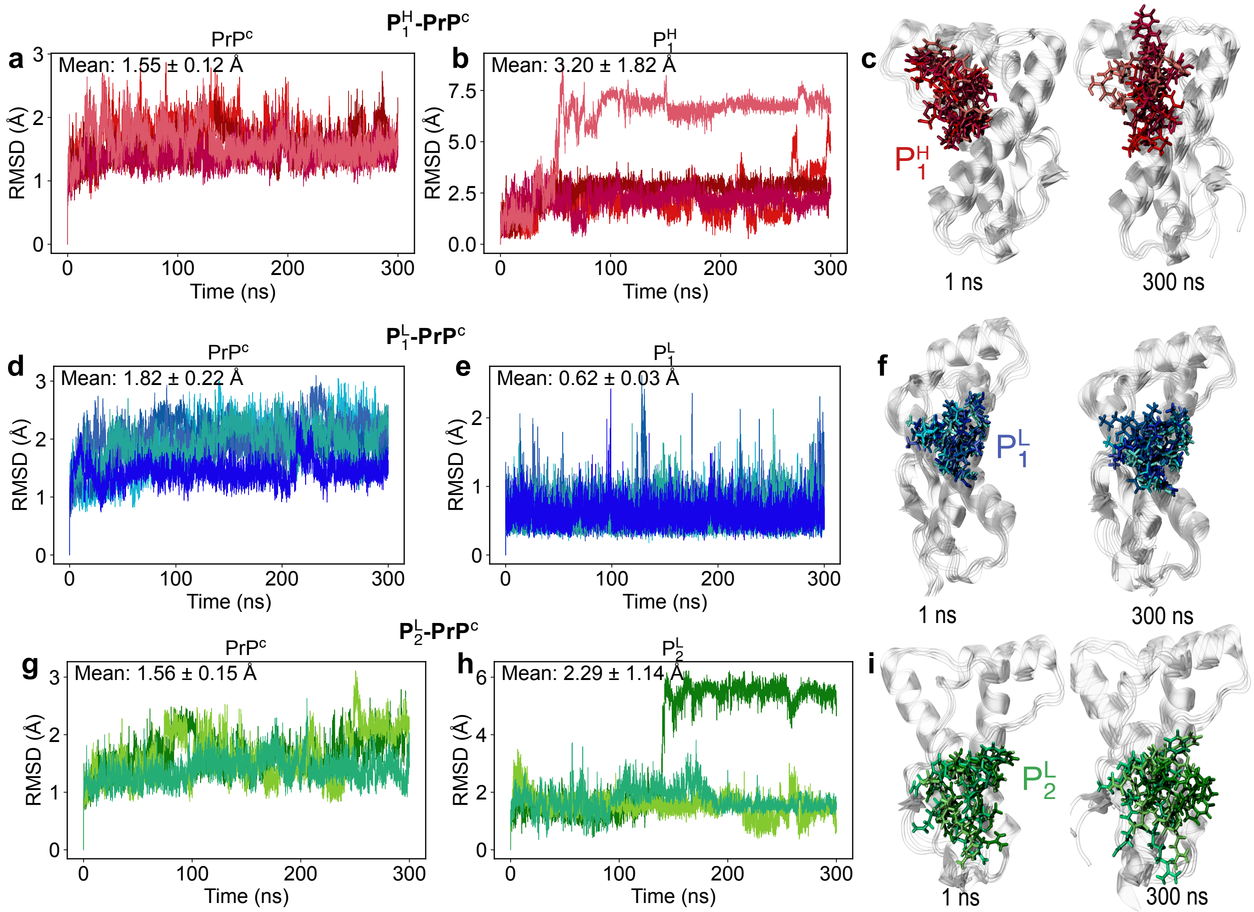


**Fig. S2 Stability of the PrP$^c$-peptide complexes.** Shown are the time series of the displacement of PrP$^c$ (a, d, g) and the peptides (b, e, h) with respect to the globular domain of PrP$^c$. The reference crystal structure is the complex with POM1 (PDB code 4H88).[1] First the structural alignment of the individual snapshots saved along the molecular dynamics simulations is carried out on the $C_\alpha$ atoms of the PrP$^c$ globular domain. Then for each simulation snapshot the peptide $C_\alpha$ root-mean-square deviation (RMSD) is calculated as $\sqrt{\frac{1}{N_{pep}}\sum_{i=1}^{N_{pep}}(\mathbf{r_i}-\mathbf{r_i^{ref}})^2}$, where $\mathbf{r_i}$ and $\mathbf{r_i^{ref}}$ are the actual and reference coordinates, respectively, of the peptide $C_\alpha$ atom $i$. $N_{pep}$ is the number of residues in the peptide. Highlighted are representative snapshots of the peptides in complex with PrP$^c$ (c, f, i).

### 3 Representative structure selection

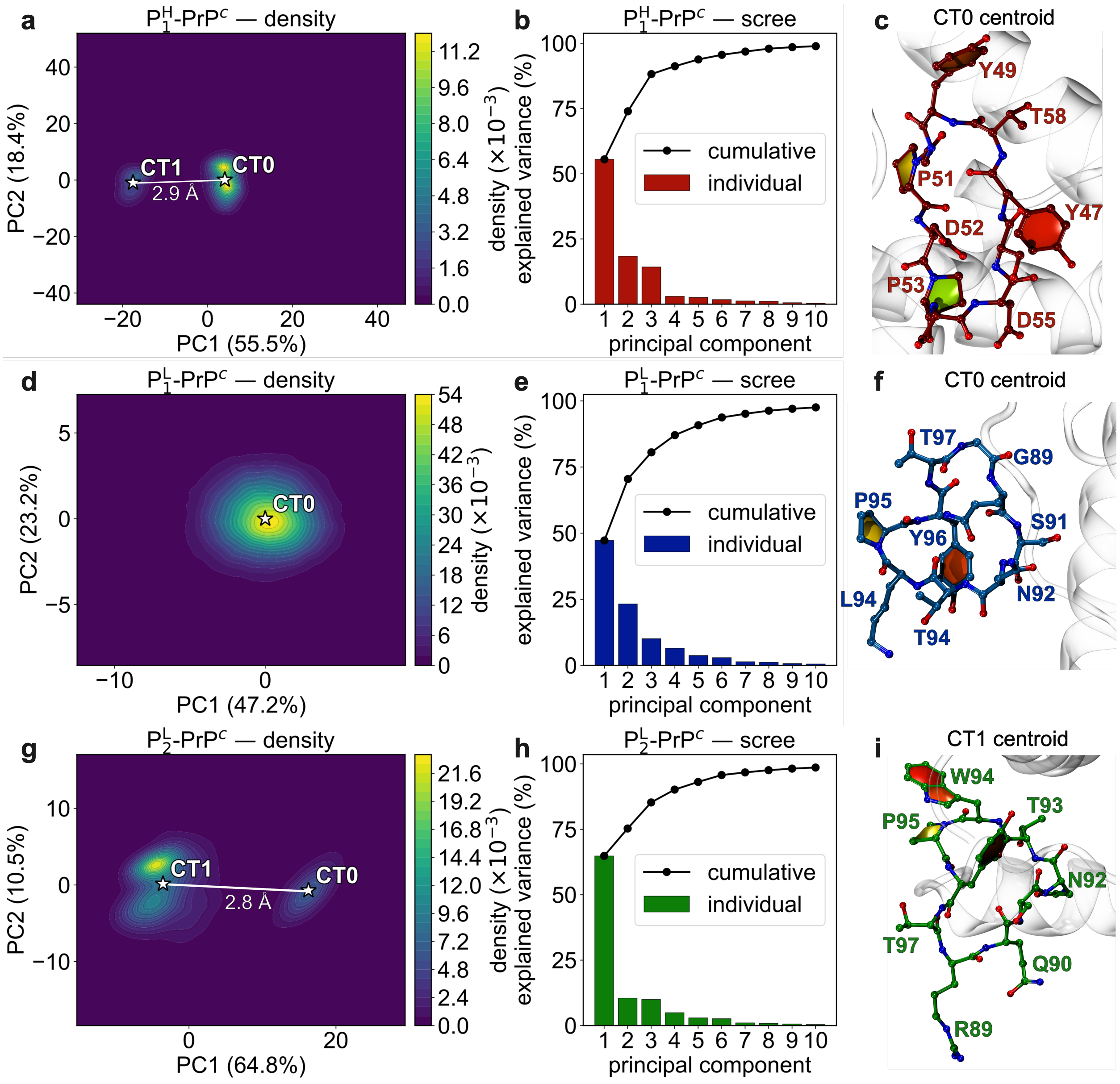


**Fig. S3 Peptide binding-pose conformational landscapes for all three PrP$^c$ peptide complexes. (a, d, g)** Kernel-density estimates of the peptide binding pose projected onto the first two principal components (PC1, PC2) for P$_1^H$-PrP$^c$ (a), P$_1^L$-PrP$^c$ (d) and P$_2^L$-PrP$^c$(g). White stars mark HDBSCAN cluster centroids (CT), and the annotated value indicates the RMSD between clusters in the PC1-PC2 plane. **(b, e, h)** Scree plots of the first ten PCs (bars: individual; line: cumulative explained variance). **(c, f, i)** Representative cluster centroids.

### 4 Western Blot analysis

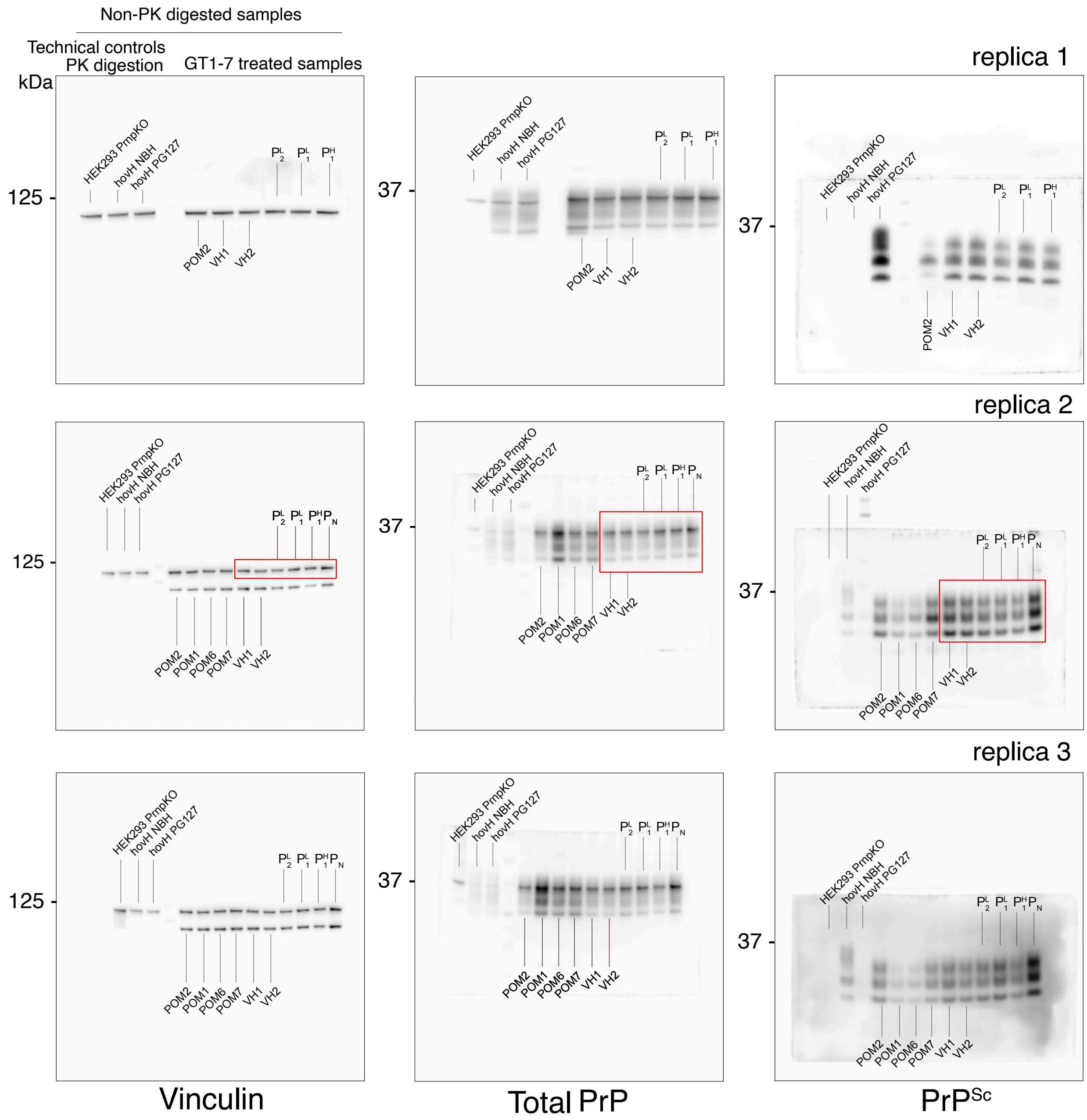


**Fig. S4 Western blot analysis.** Uncropped Western Blot images analysed for Figure 2. The regions shown in the figure are indicated by red boxes.
